# Sequence-Dependent Dynamics of U:U Mismatches in RNA Revealed by Molecular Dynamics Simulations

**DOI:** 10.1101/2025.05.14.654057

**Authors:** Miroslav Krepl, Barbora Knappeová, Jiří Šponer

## Abstract

The uracil:uracil (U:U) base pair is one of the most common mismatches observed in RNA. It is notable for its ability to adopt multiple conformational states depending on its structural environment, particularly on the identity of the flanking canonical base pairs. Here, we employed extensive molecular dynamics (MD) simulations to systematically investigate the conformational dynamics of a U:U mismatch embedded within a model A-form RNA helix, flanked by all possible canonical base pair combinations. We found that the neighboring base pairs strongly influence the preferred conformational states of the U:U mismatch. However, the mismatch still regularly samples the less favored conformations on a timescale of hundreds of nanoseconds. Contrary to previous assumptions, water-mediated conformations are not universally the most stable conformational states for isolated U:U mismatches as some of the variants distinctly prefer the direct H-bonding while others destabilize the U:U mismatch altogether. Our results strongly suggest that the presence of a U:U mismatch introduces local strain into the RNA helix, which can be relieved through dynamic destabilization of either the mismatch itself or the flanking canonical base pairs, occasionally forming a shifting “bubble” of instability. These findings advance our understanding of U:U mismatch behavior in RNA and reveal a complex interplay between local sequence context, structural stability, and RNA dynamics.

## Introduction

Deviations from the canonical Watson-Crick base pairing in A-RNA double-helices are known as base pair mismatches or non-canonical base pairs.^1–3^ The mismatches confer unique structural properties to the RNA and have significant implications for cellular function.^4^ Depending on how far their geometry differs from their canonical neighbours, the mismatches can lead to negligible changes in the helical geometry, all the way to formations of bulges or loops. The most widespread base pair mismatch in RNA is the *cis*-Watson-Crick/Watson-Crick (*c*WW) G:U wobble pair,^5–6^ which can for all intents and purposes be considered a third canonical base pair in RNA, along the *c*WW A:U and *c*WW G:C. The fourth most common *c*WW base pair in RNA is then the U:U,^3, 6–7^ which is also the most commonly occurring homo-base pair.^3, 8^ Due to its symmetry, the *c*WW U:U base pair can adopt two distinct conformational states, which differ in the relative positioning of the bases and the specific hydrogen bonds that mediate the pairing. Additionally, characteristic water insertions in the minor groove of both conformational states can occur, increasing the total number of potential conformational states of a single U:U base pair to four (Figure 1).

**Figure 1.**
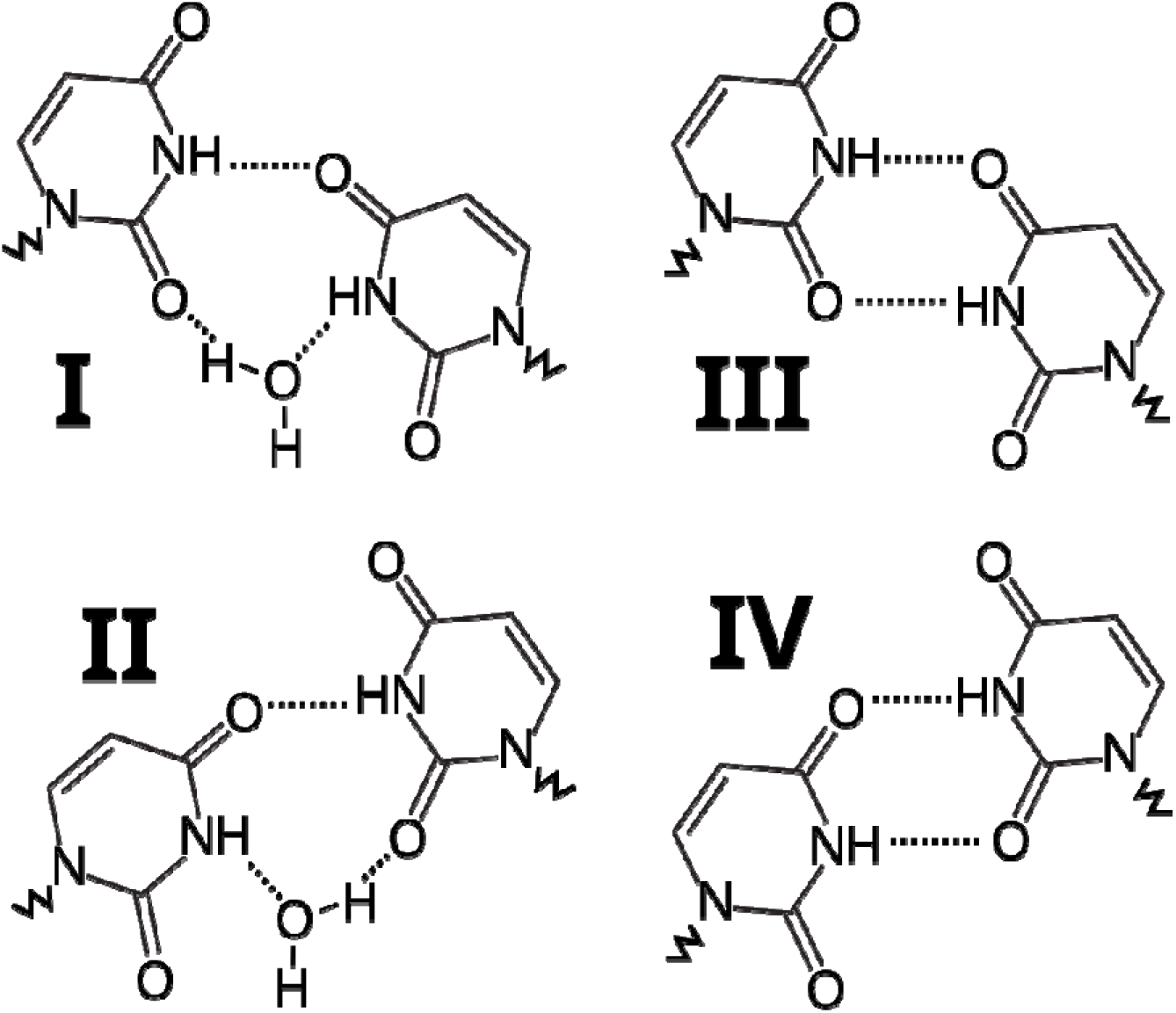
Schematic representation of the four conformational states of U:U base pairs. States I and II can be derived from states III and IV, respectively, by the insertion of a water molecule into the N3–O2 hydrogen bond. Dashed lines indicate H-bonds while jagged lines indicate the connection to the sugar-phosphate backbone (not shown). The numbering of the states follows the customs established in the literature.

In experimental structures, the cWW U:U base pair typically exhibits a preference for one of the four conformational states, depending on the specific structure and identity of the neighboring canonical base pairs. These flanking base pairs significantly influence how thermodynamically destabilizing the U:U mismatch is within the canonical A-form RNA, with some sequential contexts leading to significantly greater penalties than others.^9–10^ Although one particular U:U conformational state is seemingly always preferred in ensemble-averaged structural experiments, the potential transitions between the conformational states involve relatively minor geometrical changes. As a result, alternative conformations could transiently emerge as minor populations during thermal fluctuations, without distorting the overall helical structure. Using molecular dynamics (MD) simulations, it has indeed been shown the *c*WW U:Us undergo frequent transitions between the conformational states, giving raise to populations that can also be detected experimentally.^11–12^ The MD simulations are a computational method for studying physical movements of atoms and molecules over time, enabling a comprehensive examination of their dynamics at a spatial and temporal level currently inaccessible for experimental methods.^13^ The MD is particularly useful for studying the U:U base pair conformational transitions as these occur on timescales well accessible with the current generation of computers, providing very good convergence with no need to resort to enhanced sampling. At the same time, the U:U base pair conformational transitions occur at timescale that does not make them directly visible in the ensemble- and time-averaged structural biology experiments, further increasing the value of the computational approach.^12^

In this study, we employed molecular dynamics (MD) simulations to investigate a single U:U mismatch in all possible canonical flanking base pair contexts, represented as X<u>U</u>X trinucleotide steps, where X denotes any nucleotide and <u>U</u> is the uracil engaged in the U:U base pair. To isolate the effects of canonical base pair variation, we used a perfectly symmetrical A-RNA undecamer helix as a model system. This design ensured that any observed structural or dynamic differences can be attributed solely to the identity of the flanking base pairs. As expected, our results revealed substantial differences in the preferred U:U conformational states across different sequence contexts, while also challenging the assumptions that the water-inserted U:U conformations are uniformly the most stable.^10^ However, we also observed unexpected and significant variations in the stability of the U:U mismatch itself, and, surprisingly, in the stability of the flanking canonical base pairs. Our simulations strongly suggest that the U:U mismatch introduces structural strain into the A-form RNA helix, which can be locally alleviated by destabilizing either the U:U pair or the flanking canonical base pairs.

## Methods

### System building and simulation protocol

The starting structures for our simulations were generated with Nucleic Acid Builder (NAB) module of AMBER 22.^14^ A random palindromic undecamer with a symmetrical U:U mismatch positioned in the middle was chosen. Subsequently, all possible variants of the canonical base pairs flanking the U:U mismatch were tested. When accounting for the symmetry of the RNA helix, this corresponded to ten different systems (Figure 2). The initial files for the simulations were prepared using the xleap module of AMBER 22. The RNA was described using the OL3 *ff*.^15^ The RNA molecules were surrounded by a truncated octahedral box of OPC water molecules,^16^ ensuring a distance of 13 Å between RNA and the nearest border of the box. Potassium ions were added to obtain a bulk ion concentration of 0.15 M.^17^ Minimization and equilibration was subsequently performed following the standard RNA protocol.^11^ Afterwards, production simulations up to 10-μs-long were conducted using the Monte Carlo barostat and Langevin thermostat to maintain a pressure of 1 bar and a temperature of 300 K, respectively.^14^

**Figure 2.**
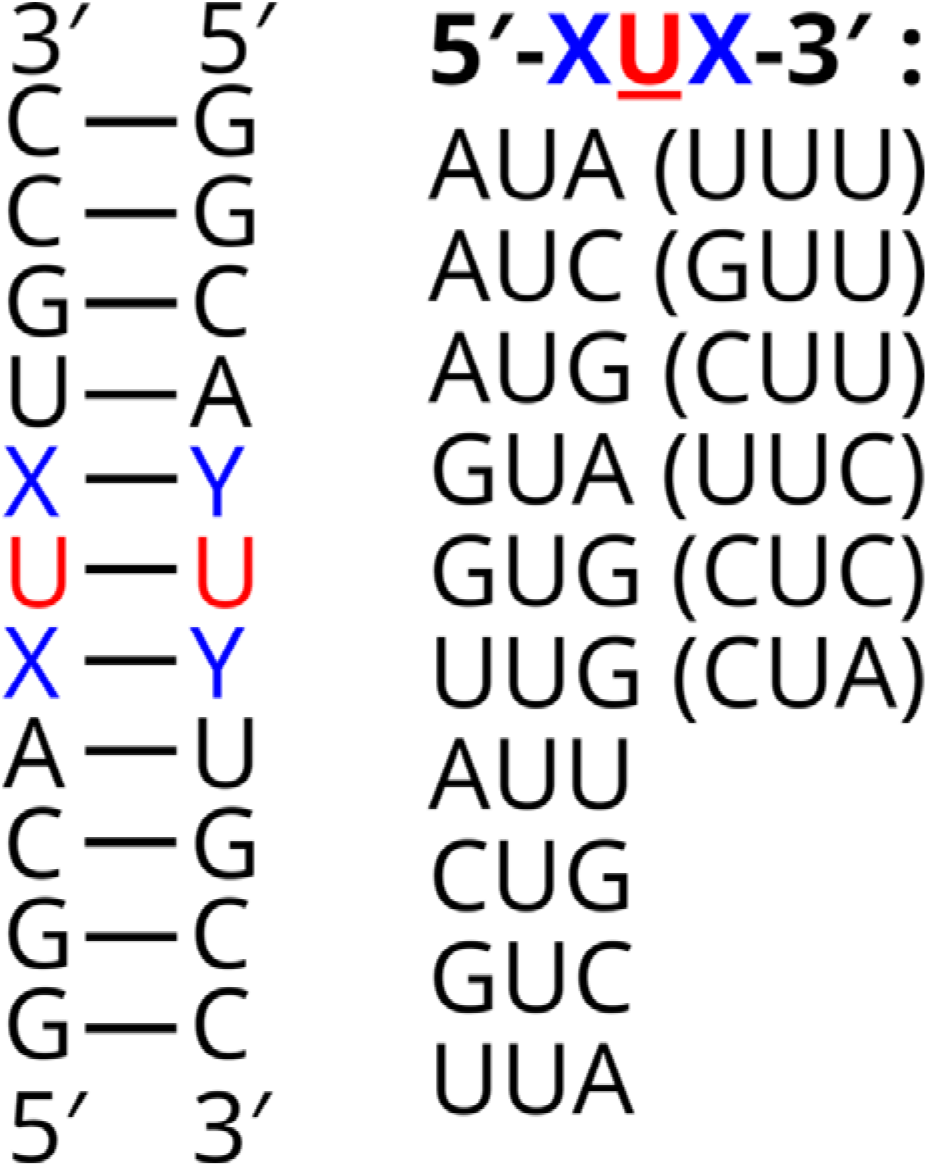
The A-RNA undecamer used in this study. The U:U mismatch is highlighted in red and its flanking canonical base pairs are shown in blue. The ten distinct variants of the “X<u>U</u>X” trinucleotide step simulated are listed on the right. Due to the molecular symmetry of the undecamer, six of the sequences (shown in parentheses) are equivalent. They are stated here to facilitate easier comparison with existing literature.

### Analyses

All trajectories were analyzed and visualized using cpptraj and VMD, respectively.^18–19^ To assess the conformational states and stability of the U:U mismatch and surrounding base pairs, we evaluated the base-pairing hydrogen bonds (H-bonds) throughout the simulations. An individual H-bond was considered present if the donor–acceptor distance was less than 4 Å and the donor–hydrogen–acceptor angle exceeded 120°. The starting structures were modeled as standard A-RNA helix and the U:U base pair was not initially formed properly. To account for this, the first 250 ns of all trajectories were considered as extended equilibration period and not included in the analyses. Unless stated otherwise, the results below will be referring to the ten-microsecond simulations.

## Results and Discussion

In this section, we explore the conformational dynamics of U:U base pairs, considering all the possible variations in its immediately neighboring (flanking) canonical base pairs, while keeping the rest of the sequence unchanged and symmetrical. Surprisingly, each trinucleotide step containing the single U:U base pair shows distinct behaviour. Namely, there are differences in preferred base pair conformations, balance between direct and water-mediated H-bonding, and stability of the U:U base pair and its flanking canonical base pairs. The total number of simulations was 50 with cumulative length of 260 μs (Table 1).

**Table 1.** List of simulations.

| System<sup>a</sup> | number of simulations × length [μs] |
| --- | --- |
| 5'-AUA-3' | 2 × 10, 3 × 2 |
| 5'-AUC-3' | 2 × 10, 3 × 2 |
| 5'-AUG-3' | 2 × 10, 3 × 2 |
| 5'-AUU-3' | 2 × 10, 3 × 2 |
| 5'-CUG-3' | 2 × 10, 3 × 2 |
| 5'-GUA-3' | 2 × 10, 3 × 2 |
| 5'-GUC-3' | 2 × 10, 3 × 2 |
| 5'-GUG-3' | 2 × 10, 3 × 2 |
| 5'-UUA-3' | 2 × 10, 3 × 2 |
| 5'-UUG-3' | 2 × 10, 3 × 2 |

<sup>a</sup>For clarity, the uridine which is part of the U:U mismatch is underlined and presented as such throughout the paper. Note that mathematically the number of possible variations is sixteen, however, due to molecular symmetry of the undecamer, only ten unique variants exist (see also Figure 2).

### Conformational preference of the U:U base pairs

Out of the ten simulated systems, four of them (G<u>U</u>C, C<u>U</u>G, U<u>U</u>A, A<u>U</u>U) have the flanking canonical base pairs forming a perfectly symmetrical environment. In other words, rotating such helix by 180° does not alter the displayed sequence (Figure 2). Because of this symmetry, and assuming a sufficient convergence of the MD simulations, these four systems should give exactly equal populations of the III(I) v. IV(II) U:U base pair states (Figure 1). Indeed, in all our ten-microsecond simulations we observed the ratio between the III(I) and IV(II) states close to ~1:1 for all four systems. We also observed a good match between the two independent trajectories (Table 2). This indicates a decent, albeit not perfect convergence can be achieved for the U:U base pair conformational dynamics on the ten-microsecond timescale using standard MD simulations. Notably, the convergence was clearly not yet sufficient on two-microsecond timescale, where we observed large deviations from the expected 1:1 ratio for the four symmetrical systems, as well as relatively large differences between some of the independently run trajectories of the same systems (Supporting Information Table S1).

**Table 2.**
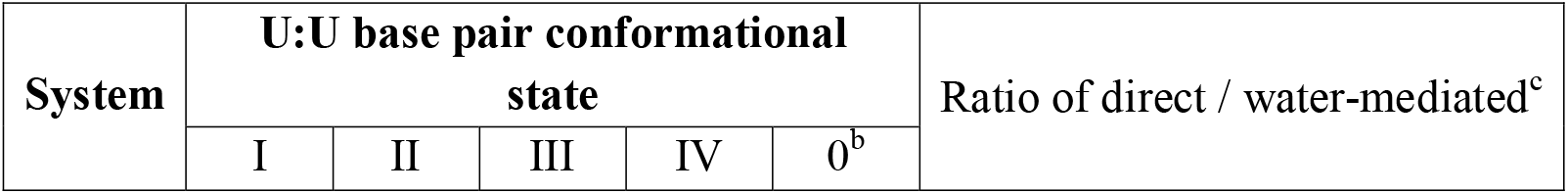
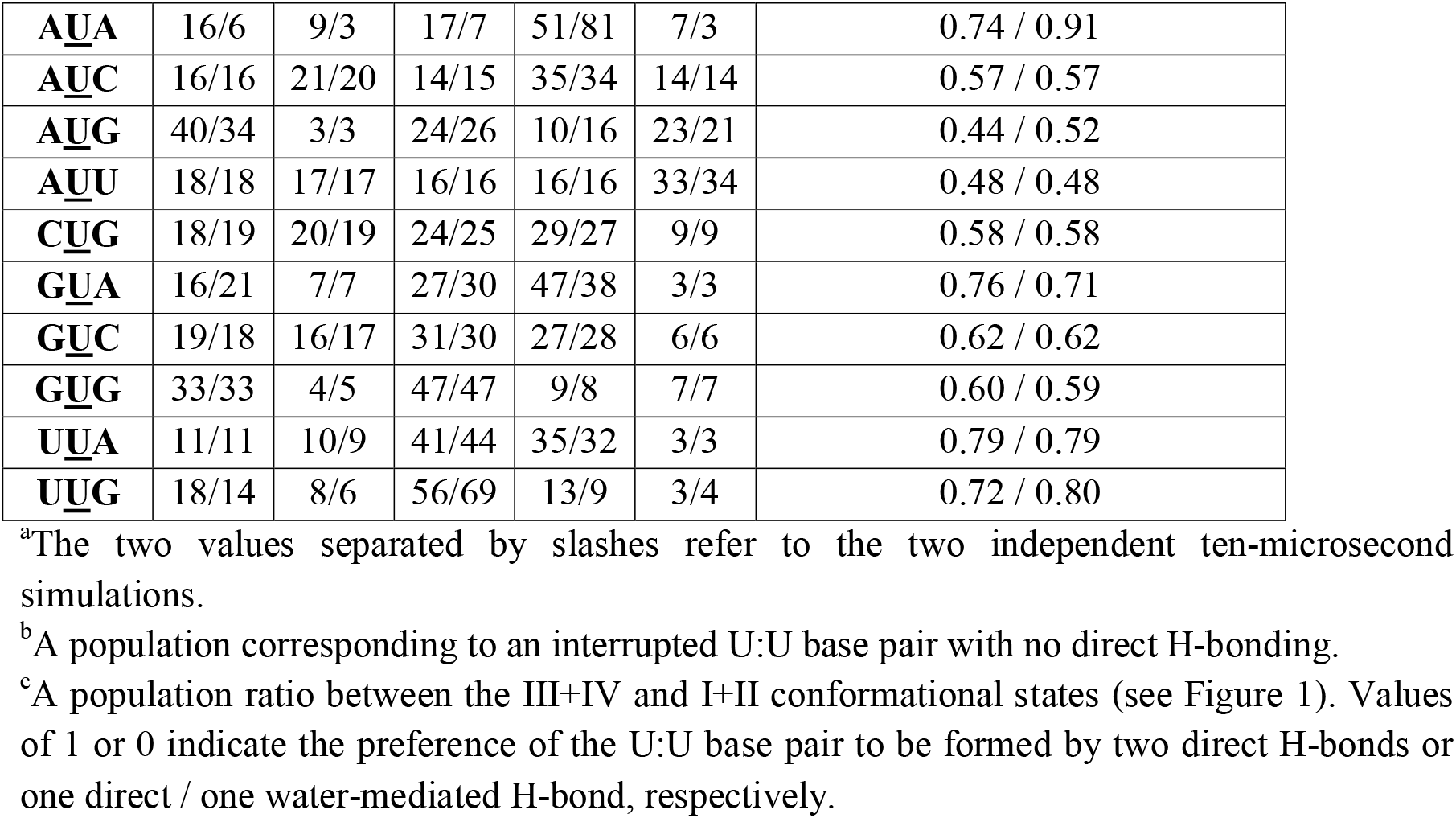
Populations of the U:U base pair conformational states (in %).

In rest of the systems, the preferred U:U state in simulations was III(I) and IV(II) for sequences A<u>U</u>G, G<u>U</u>G, U<u>U</u>G and A<u>U</u>A, A<u>U</u>C, G<u>U</u>A, respectively. Strongest respective preferences were observed for the G<u>U</u>G and A<u>U</u>A systems (Table 2). The frequency of the transitions and their lifetimes varied across the systems, but generally occurred on the scale of tens to hundreds of nanoseconds (Figure 3). Importantly, the U:U transitions occurred constantly throughout all the simulations with approximately constant frequency, with no signs of the systems “settling” into one specific conformational state.

**Figure 3.**
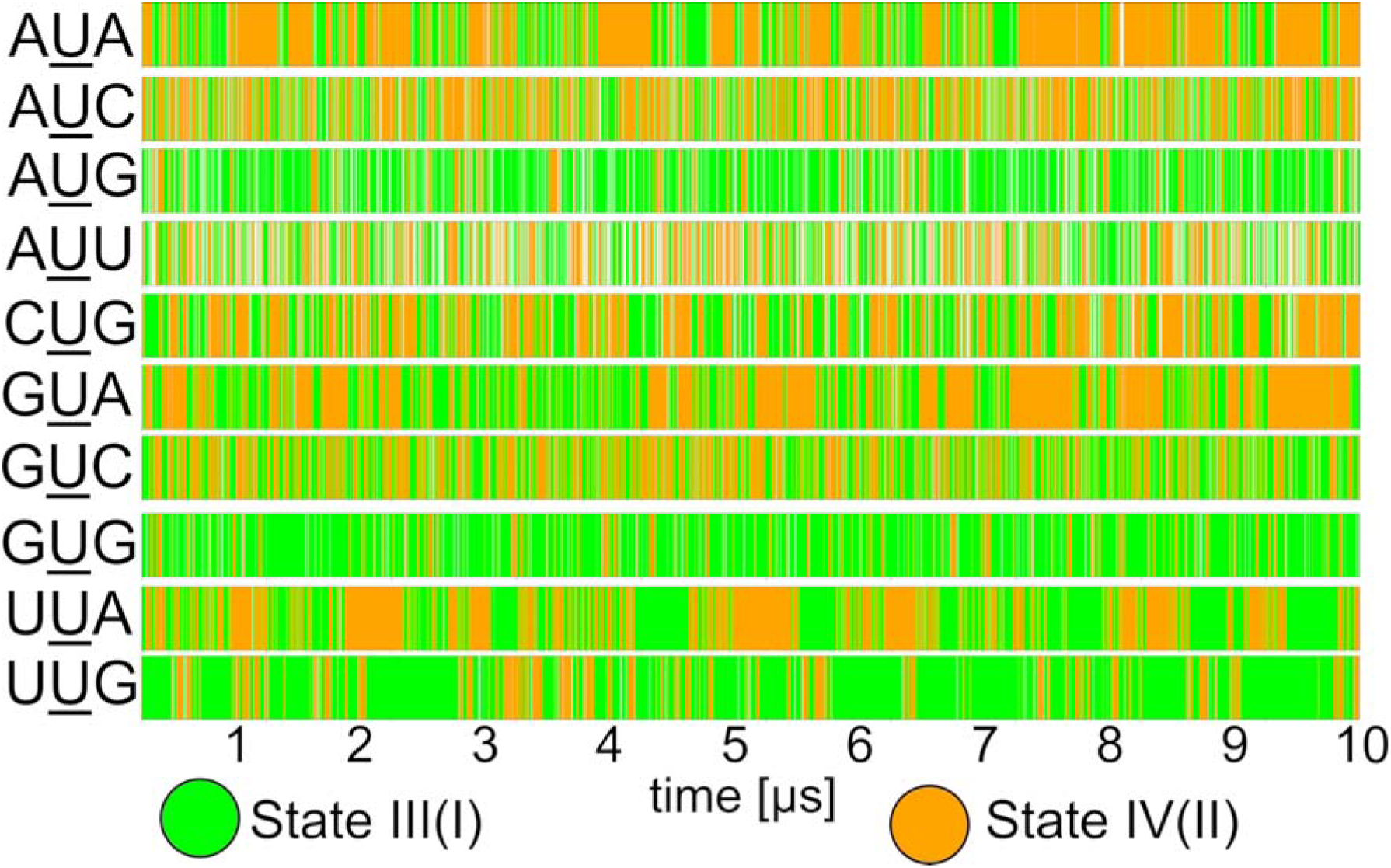
Conformational transitions and reversible interruptions of the U:U base pair in MD simulations. Time evolution of the U:U mismatch in a selected 10-μs simulation for each system. See Figure 1 for the definition of the conformational states. White segments represent reversible interruptions of the U:U during which no base-pairing H-bond was present (mostly A<u>U</u>U system; see also Supporting Information Figure S1). Note that due to their brief lifetimes and rapid exchange rates, the partially water-mediated states I and II cannot be effectively differentiated in these plots.

### The U:U base pairs exhibit characteristic balance between direct and water-mediated H-bonding and complete instability

In addition to the balance between the U:U conformational states (see above), the identity of the flanking canonical base pairs also influenced whether two direct H-bonds (states III and IV; 2 H-bond) or one direct and one water-mediated H-bond are preferred (states I and II; 1 H-bond). Most systems preferred the 2 H-bond states III/IV slightly more, with the notable exception of the A<u>U</u>G and A<u>U</u>U systems where the water-mediated states I/II dominated (Table 2). We note the transitions between 1 H-bond and 2 H-bond states also involve a change in the C1′-C1′ distances of the U:U base pair. In fact, for 1 H-bond states, this distance closely corresponded to the standard C1′-C1′ distances observed for the canonical A-RNA base pairs (~10.4 Å). The preference for either 2 H-bond or 1 H-bond arrangement of U:U can therefore also be interpreted in terms of how strongly the flanking canonical base pairs enforce values typical for A-RNA helix (Figure 4).

**Figure 4.**
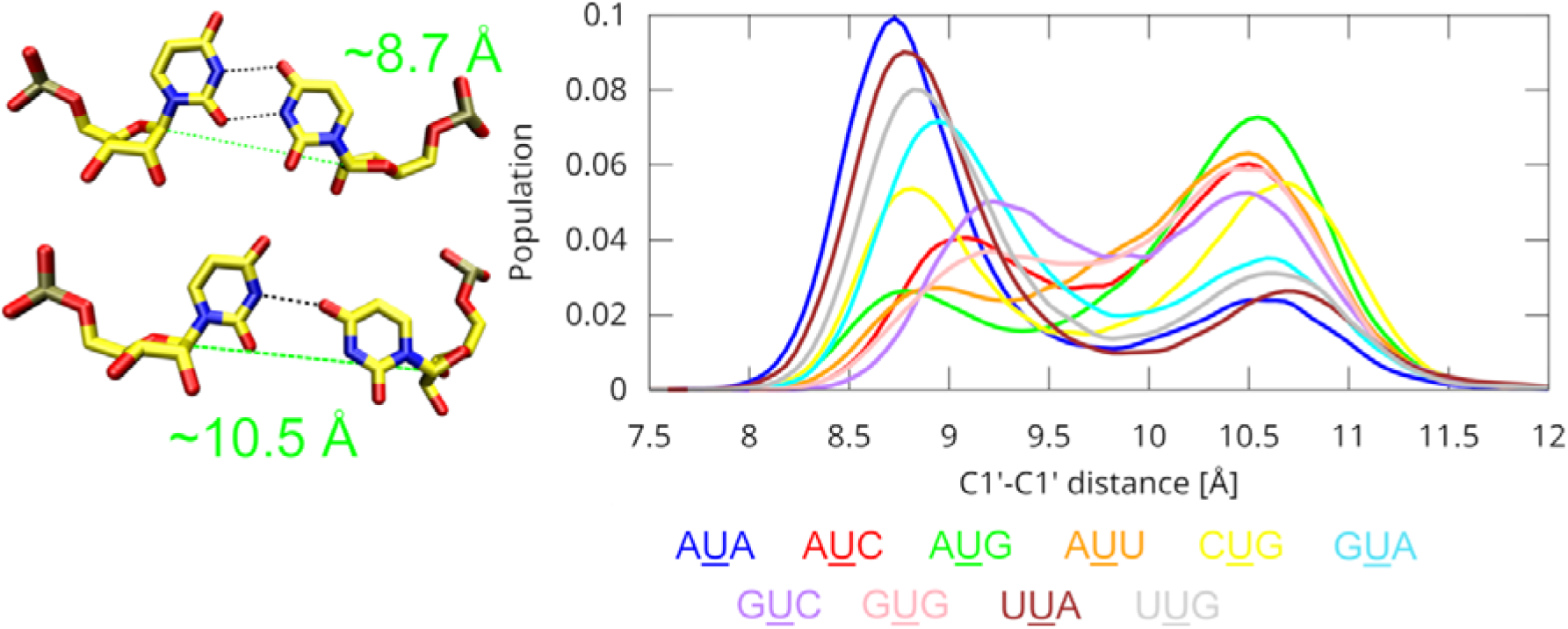
C1′–C1′ distances of the U:U base pair. A significantly larger C1′–C1′ distance (green dashed line), matching the typical value observed in canonical A-form RNA, is found when only a single direct base-pairing H-bond (black dashed lines) is present. The histogram on the right displays the distribution of C1′–C1′ distances for the U:U base pair in combined simulation ensembles of the two ten-microsecond simulations. Individual systems are color-coded according to the legend shown below. In addition to the different distribution, notable differences between systems are evident also in the positions of the peak maxima, indicating variation in the preferred C1′–C1′ distances across different sequence contexts.

In addition to the 2 H-bond and 1 H-bond states, we also observed instances when no direct H-bonding existed between the U:U base pair at all (Table 2). While minor (under ~10%) in most systems, this state was prominent for e.g. the A<u>U</u>U system, where it populated ~33% of the simulation ensemble (Figure 3). We note that the C1′-C1′ distances of this “0 H-bond” state were almost exact match for the standard canonical base pairs. It can therefore be considered a more severe response to the same effects of the flanking canonical base pairs that promote the 1 H-bond state for U:U. In fact, sequences which displayed greater affinity for the 1 H-bond state over the 2 H-bond state also had the most significant population of the 0 H-bond state (Table 2). Structurally, the 0 H-bond state is characterized by the uridine bases drifting apart beyond the typical H-bonding distance (Supporting Information Figure S1).

Notably, no bulging or larger structural distortions associated with the U:U base pair disruptions were observed at any point, and they remained fully reversible on our simulation timescale.

### Instability of the flanking canonical base pairs

In five of the systems (A<u>U</u>**A, C**<u>U</u>**G**, G<u>U</u>**A, U**<u>U</u>**A, U**<u>U</u>G; the bold indicates the unstable base pair), we observed reversible instability of one or both of the flanking canonical base pairs. There was an anticorrelation between the instability of the U:U base pair (Table 2) and its flanking canonical base pairs (Table 3). In other words, the systems tended to either have increased population of the 0 H-bond state of the U:U (see above) or unstable flanking canonical base pairs, but not both, suggesting a structural struggle between them. A particularly striking example is provided by the AUU and UUA systems, which differ only in the polarity of the trinucleotide step, yet show pronounced instability in the U:U mismatch and flanking base pairs, respectively. We note that the flanking canonical base pair displaying the reversible instability was always A:U/U:A, reflecting their lower thermodynamic stability compared to G:C/C:G. The notable exception was the C<u>U</u>G which displayed both the flanking canonical base pair instability as well as modest population of the 0 H-bond state of the U:U. Interestingly, earlier thermodynamic experiments indicated the U:U mismatch causing the greatest destabilization of the helix in the C<u>U</u>G sequential context.^9^ We suggest our data rationalizes this effect. We also observe that the stability of the flanking base pairs is strongly influenced by the conformational state of the U:U mismatch. Specifically, U:U conformational states III(I) and IV(II) tend to destabilize the preceding and succeeding canonical base pairs, respectively (Figure 3 and Figure 5). This effect reflects the greater stacking overlap between the U:U mismatch and one of the neighboring base pairs in each conformational state. Lastly, we note that no similar base pair disruptions were observed among the rest of the base pairs within the helix. The instability thus appears to be fully contained to the immediate vicinity of the U:U mismatch, at least on our simulation timescale.

**Table 3.** Stability (in %) of the canonical base pairs flanking the U:U mismatch base pair.^a^.

| System<sup>a</sup> | 5' | 3' |
| --- | --- | --- |
| **AUA** | 98/99 | 62/59 |
| **CUG** | 91/90 | 89/92 |
| **GUA** | 100/100 | 62/71 |
| **UUA** | 75/70 | 78/82 |
| **UUG** | 62/41 | 97/97 |

<sup>a</sup>The 5' and 3' indicate values for the flanking canonical base pairs located upstream and downstream of the mismatch uridine, respectively. The remaining systems exhibited no significant instability in the flanking canonical base pairs beyond regular thermal fluctuations (data not shown). Note that populations of ~97-98% should be considered as fully stable canonical base pairing under our analyses criteria, accounting for expected thermal fluctuations and the defined cutoffs (see Methods).

**Figure 5.**
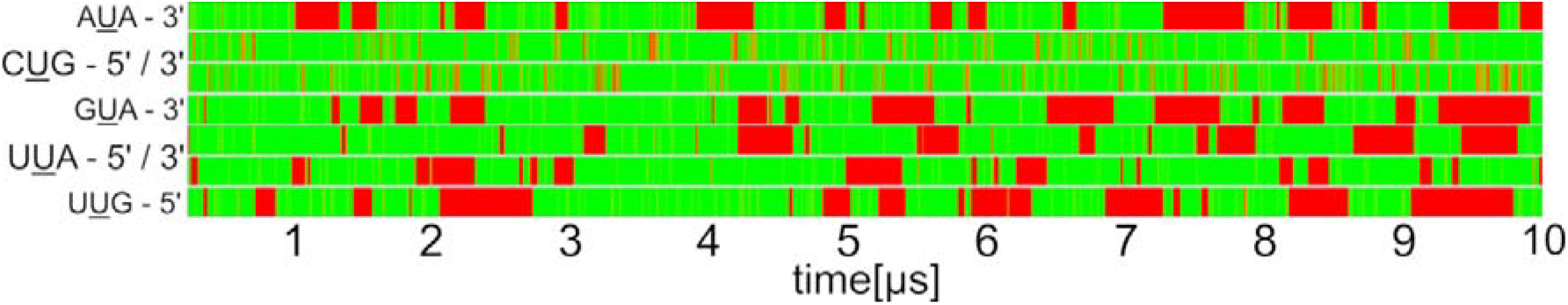
Reversible interruptions of the canonical base pairs flanking the U:U mismatch. Green and red indicate stable and interrupted base pairs, respectively. A base pair was classified as interrupted whenever none of the H-bonds defining a standard *c*WW base pair were present. For systems where the values for both the 5′ and 3′ flanking canonical base pairs are stated, the 5′ is the one on top.

## Conclusions

In this work, we employed molecular dynamics (MD) simulations to investigate the dynamics of the U:U mismatch base pair in all possible nucleotide contexts defined by its flanking canonical base pairs. Our findings challenge the prevailing view that the flanking sequence merely determines the conformational state of the U:U mismatch or that U:U mismatches inherently favor water-mediated conformations. While these trends are generally observed, at least for some of the sequences, our study reveals a far more complex interplay of energetic contributions within the local structural environment. A key observation is that the presence of the U:U mismatch introduces structural strain, which can be alleviated in two principal ways: either by destabilizing the U:U pair to allow it to adopt a quasi-canonical A-RNA geometry, or by destabilizing one or both of the flanking canonical base pairs to better accommodate the narrower U:U mismatch. Importantly, this balance is highly delicate, leading to frequent conformational exchanges on the timescale of our simulations. In some sequences, this takes form of a transient “bubble” of instability that shifts dynamically along the trinucleotide region of the U:U mismatch. Overall, our data provide a comprehensive overview of how flanking canonical sequences influence the dynamics and stability of U:U base pairs. These insights can guide the rational design of functional RNA molecules containing such mismatches and aid in the interpretation of biochemical data.

## Supporting Information

Supporting Tables and Figures.

## Acknowledgement

This work has been conducted in the sustainability period of the project SYMBIT No. CZ.02.1.01/0.0/0.0/15_003/0000477 as its follow-up activity. We acknowledge the use of CESNET data storage facilities [grant number LM2018140].

