## Supporting Information for "Sequence-Dependent Dynamics of U:U Mismatches in RNA Revealed by Molecular Dynamics Simulations"

### Supporting Information Tables

Table S1. Populations of the U:U base pair conformational states (in %).

| **System** | **U:U base pair conformational state** | | | | |
| --- | --- | --- | --- | --- | --- |
|  | I | II | III | IV | 0^b^ |
| **AUA** | 7/16/16 | 7/6/9 | 7/16/15 | 75/53/53 | 4/8/7 |
| **AUC** | 18/16/15 | 19/19/21 | 16/16/15 | 32/35/35 | 15/14/15 |
| **AUG** | 38/38/38 | 4/3/4 | 22/25/23 | 13/11/14 | 23/24/22 |
| **AUU** | 15/16/18 | 19/17/17 | 14/16/16 | 19/16/15 | 34/35/34 |
| **CUG** | 9/14/20 | 20/22/20 | 17/19/23 | 42/30/26 | 12/16/11 |
| **GUA** | 9/15/22 | 8/7/9 | 12/21/38 | 70/54/27 | 2/3/3 |
| **GUC** | 17/18/17 | 20/18/19 | 28/29/28 | 30/31/31 | 5/6/5 |
| **GUG** | 33/37/34 | 6/3/4 | 45/49/47 | 10/4/8 | 6/6/7 |
| **UUA** | 10/12/5 | 14/12/12 | 26/35/14 | 45/37/66 | 4/4/2 |
| **UUG** | 16/21/24 | 7/6/11 | 62/59/42 | 12/11/18 | 3/4/5 |

^a^The three values separated by slashes refer to the three independent two-microsecond simulations.

^b^A population corresponding to an interrupted U:U base pair.

**Supporting Information Figures**

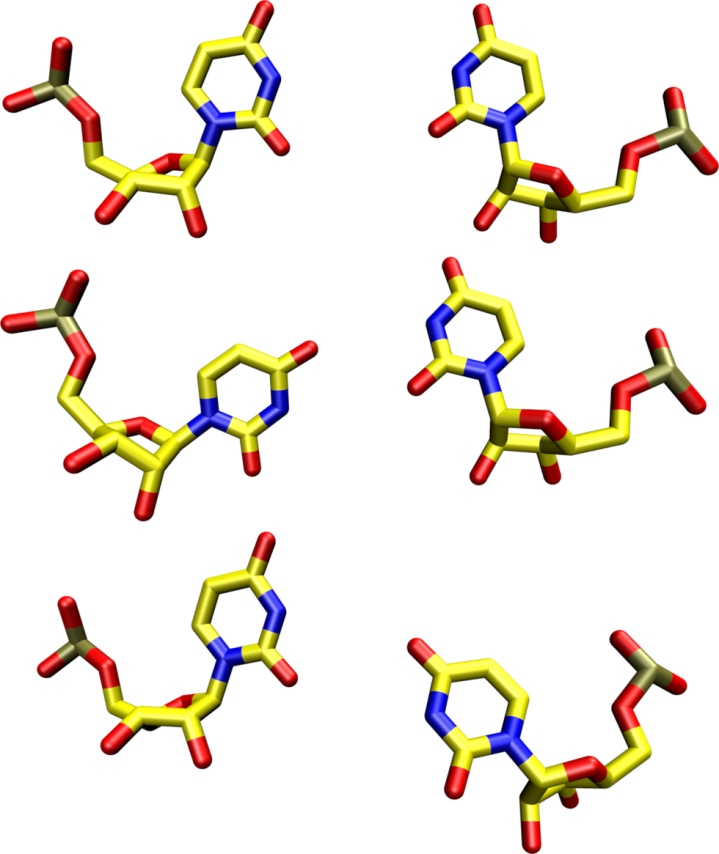

Figure S1. **The “0 H-bond” state of the U:U base pair.** The three base pairs displayed are the arrangements that were regularly observed in the MD simulations when the U:U base pair reversibly disintegrated. In each of them, the C1′–C1′ distance corresponded to the canonical A-RNA and there were no direct H-bonds between the uridines, with the space between them occupied by water molecules. Importantly, we did not observe any bulging or other large scale distortions.
